# The conserved histone chaperone LIN-53 links lifespan and healthspan regulation in *Caenorhabditis elegans*

**DOI:** 10.1101/539015

**Authors:** Stefanie Müthel, Bora Uyar, Mei He, Anne Krause, Burcu Vitrinel, Selman Bulut, Djordje Vasiljevic, Altuna Akalin, Stefan Kempa, Baris Tursun

## Abstract

Whether extension of lifespan provides an extended time without health deteriorations is an important issue for human aging. However, to which degree lifespan and healthspan regulation might be linked is not well understood. Chromatin factors could be involved in linking both aging aspects, as epigenetic mechanisms bridge regulation of different biological processes. The epigenetic factor LIN-53 (RBBP4/7) is required for safeguarding cell identities in *Caenorhabditis elegans* as well as mammals and for preventing memory loss and premature aging in humans. LIN-53 is a histone chaperone that associates with different chromatin-regulating complexes. We show that LIN-53 interacts with the Nucleosome remodeling and deacteylase (NuRD)-complex in *C. elegans* muscles to promote healthy locomotion during aging. While mutants for other NuRD members show a normal lifespan, animals lacking LIN-53 die early because LIN-53 depletion affects also the Histone deacetylase complex Sin3, which is required for a normal lifespan. To determine why *lin-53* and *sin-3* mutants die early, we performed transcriptome and metabolome analysis and found that levels of the disaccharide Trehalose are significantly decreased in both mutants. As Trehalose is required for normal lifespan in *C. elegans, lin-53* and *sin-3* mutants could be rescued by either feeding with Trehalose or increasing Trehalose levels via the Insulin/IGF1 signaling pathway. Overall, our findings suggest that LIN-53 is required for maintaining lifespan and promoting healthspan through discrete chromatin regulatory mechanisms. Since both LIN-53 and its mammalian homologs safeguard cell identities, it is conceivable that its implication in lifespan and healthspan regulation is also evolutionarily conserved.

## Introduction

The decline of physical condition and the onset of diseases such as cancer, diabetes or dementia are important issues during aging. Age-associated deterioration of health has gained importance as the human life expectancy constantly increases worldwide. It has been predicted that in 2050 adults over the age of 80 will triple compared to the year 2015 (Jaul & Barron, 2017). Hence, an important aspect of aging is whether increasing lifespan would also extend the healthspan, meaning the time of life without unfavorable health conditions. However, genetic factors that play a role in linking healthspan with lifespan regulation are largely unknown. Aging regulation by chromatin-regulating factors could play a role in linking lifespan with healthspan as loss of epigenetic gene regulation diminishes cell fate safeguarding (Kolundzic et al., 2018; Onder et al., 2012; Yadav, Quivy, & Almouzni, 2018), declines stem cell health (Brunet & Rando, 2017; Ren, Ocampo, Liu, & Belmonte, 2017), impairs muscle regeneration (Guasconi & Puri, 2009) and shortens lifespan of organisms (Field & Adams, 2017; Greer et al., 2010).

One specific type of epigenetic regulators are histone-chaperones, which are proteins that directly interact with histones and function as a scaffold for chromatin-modifying protein complexes (Hammond, Strømme, Huang, Patel, & Groth, 2017). They are important for folding, oligomerization, post-translational modifications, nucleosome assembly, and genomic location of histones (Hammond et al., 2017). The *C. elegans* histone-chaperone LIN-53 is highly conserved and known as RBBP4/7 (also as CAF-1p48) in mammals. LIN-53 and its homologs can be found in different protein complexes that regulate the repressive and active state of chromatin (Lu & Horvitz, 1998) (Loyola & Almouzni, 2004) (Eitoku, Sato, Senda, & Horikoshi, 2008). Among those complexes are PRC2 (Polycomb repressive complex 2, (Margueron & Reinberg, 2011)), Sin3 Histone Deacetylase Complex (Sin3 HDAC) (Nicolas et al., 2000), NuRD (<u>Nu</u>cleosome <u>r</u>emodelling and <u>d</u>eacetylase complex (Allen, Wade, & Kutateladze, 2013)), CAF-1 histone-chaperone complex (Verreault, Kaufman, Kobayashi, & Stillman, 1996) and DRM (Dp/Rb/Muv (Harrison, Ceol, Lu, & Horvitz, 2006)). In *C. elegans,* LIN-53 was shown to interact with the Rb homolog LIN-35 to antagonize the Ras signaling pathway (Lu & Horvitz, 1998). Moreover, LIN-53 and its mammalian homologs RBBP4/7 safeguard cells against reprogramming (Cheloufi et al., 2015; Tursun, Patel, Kratsios, & Hobert, 2011) and have been implicated in age-related memory loss and premature aging in humans (Pavlopoulos et al., 2013) (Pegoraro et al., 2009).

In this study, we revealed that LIN-53 is required for healthy motility and normal lifespan in *C. elegans*. Notably, the muscle defects and the short lifespan in *lin-53* mutants can be unlinked based on different chromatin-regulating complexes. LIN-53 is interacting with the NuRD complex to maintain muscle integrity and proper motility but requires the Sin3 complex to ensure normal lifespan. To understand why *lin-53* and *sin-3* mutants have a shortened lifespan, we analyzed the transcriptome as well as metabolome of mutant animals. Loss of LIN-53 or SIN-3 leads to a strong decrease in Trehalose levels - a disaccharide that is required for a normal lifespan (Y. Honda, Tanaka, & Honda, 2010), (Seo, Kingsley, Walker, Mondoux, & Tissenbaum, 2018). Restoring Trehalose levels by feeding, or genetically via the Insulin/IGF1 signaling (IIS) pathway, suppressed the short lifespan of *lin-53* and *sin-3* mutants, supporting the idea that LIN-53 and SIN-3 are required to maintain a normal lifespan via ensuring the homeostasis of metabolites such as Trehalose.

Overall, our findings suggest that the epigenetic factor LIN-53 links healthspan and lifespan regulation in *C. elegans*. As LIN-53 is a highly conserved chromatin regulator with an evolutionarily conserved role in cell fate safeguarding (Cheloufi et al., 2015; Tursun et al., 2011), (Cheloufi & Hochedlinger, 2017), it is conceivable that its homologs regulate lifespan and healthspan also in other species. Hence, our findings provide an initial framework for elucidating how lifespan and healthspan regulation might be linked through epigenetic factors, which could be of high relevance for human health and aging.

## Results

### Loss of LIN-53 results in muscle and locomotion defects

The role of the highly conserved histone chaperone LIN-53 in somatic tissues of *C. elegans* is poorly understood. We therefore examined *lin-53* null mutants and noticed a severe movement defect for *lin-53(n3368)* animals. They exhibit decreased mobility on solid agar plates (Fig. 1A) as well as in liquid when compared to wild-type animals (Fig. 1B) at the larval L4 stage, young adult stage and as 2 day old adults. Such motility defects can point to an impaired muscle apparatus, which prompted us to stain muscles and assess their integrity in *lin-53* mutants. Using fluorescent Phalloidin, which binds to F-actin fibers in muscles (Fig. 1C), and an antibody against the myosin-heavy chain (MHC) component of body wall muscles (Fig. 1D), we observed disrupted muscle structures in *lin-53* mutants (Fig. 1C and 1D). The decline in muscle integrity upon *lin-53* depletion is also evident based on animals expressing a *Pmyo-3::GFP* reporter (Fig. 1E). These muscle phenotypes in *lin-53* mutants are cell-autonomous effects as muscle-specific RNAi against *lin-53,* by using a hairpin construct (*myo-3p::lin-53* HP) also leads to muscle and motility defects (Fig. S1A and S1B). Consequently, the muscle and motility defects in *lin-53* mutants can be rescued by expressing full length LIN-53 specifically in muscles using the *myo-3* promoter (Fig. 1F and Fig. S1C). Interestingly, wt animals overexpressing *myo-3p::lin-53* or *myo-3p::GFP::lin-53* move significantly better than control animals suggesting that the overexpression of LIN-53 in muscles has a beneficial effect to maintain the motility in adult animals (Fig. S1D).

**Fig. 1.**
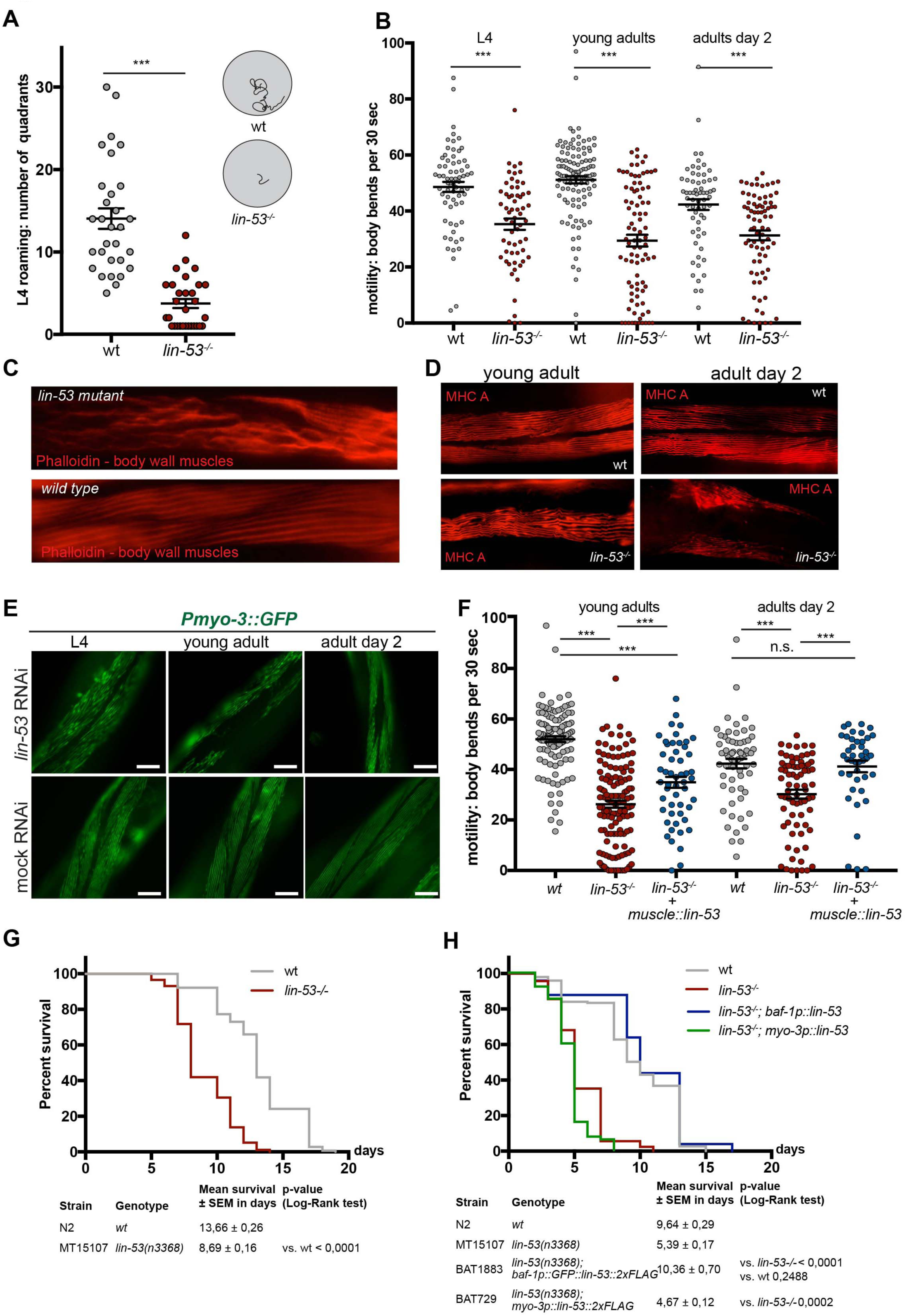
Loss of LIN-53 causes locomotion defects and short lifespan. (A) L4 control and *lin-53* mutant worms were put on solid agar plates and the movement on solid agar was monitored using graph paper (sketch on the right side of the graph). *lin-53* mutant animals move significantly slower than control animals. Statistical analysis was carried out using unpaired t-test; *** p < 0,0001. (B) Control and *lin-53* mutant animals at three different developmental stages (L4 larvae, young adult animals and adults at day 2) were put in M9 medium, motility was recorded and the body bends were counted using the ImageJ WrmTrck plugin. At all developmental stages *lin-53* mutants swim significantly slower than control animals. Statistical analysis was carried out using unpaired one-way ANOVA; *** p < 0,0001. (C) Phalloidin binds to F-actin in body wall muscles. In *lin-53* mutants animals the muscle structure is disrupted compared to control animals. (D) The muscle structure of *lin-53* mutants and control animals at two different developmental stages (young adult and adult day 2) was analyzed using an antibody against MHC in immunostaining. The muscle structure is disrupted in *lin-53* mutants compared to control animals at both stages. (E) Worms expressing a *Pmyo-3::GFP* reporter were subjected to control and *lin-53* RNAi at different developmental stages (L4, young adults, adult day 2). At all three stages a muscle structure disruption is detectable upon loss of *lin-53.* (F) Expression of recombinant LIN-53 only in muscles rescues the motility defect of *lin-53* mutants. Statistical analysis was carried out using unpaired one-way ANOVA; *** p < 0,0001, ns = not significant. (G) Depletion of *lin-53* decreases the lifespan of *C. elegans* by 5 days (p-value < 0,0001). wt animals (grey line; mean lifespan 13,66 ± 0,26 days), *lin-53* mutants (red line; mean lifespan 8,69 ± 0,16 days). Triplicate experiments with 40 animals per repeat. Survival analysis was carried using Kaplan-Meier-estimator, p-value was calculated using Log-Rank Test. (H) The short lifespan of *lin-53* mutants is not rescued upon overexpression of *lin-53* in muscles but upon ubiquitous expression using the *baf-1* promoter. Triplicate experiments with 40 animals per repeat.Survival analysis was done using Kaplan-Meier-estimator, P-value was calculated using Log-Rank Test.

Overall, our findings suggest that LIN-53 is required to maintain muscle integrity and to prevent the decline of locomotion capabilities in *C. elegans*.

### Loss of LIN-53 shortens lifespan

Since deterioration of coordinated movement is associated with aging (Herndon et al., 2002), we wondered whether *lin-53* mutants suffer from a short lifespan. Lifespan assays revealed that *lin-53(n3368)* mutants have an average lifespan which is around 40% shorter than wt animals (Fig. 1G and Table S1). A shortened lifespan is also evident in animals carrying the CRISPR/Cas9-generated *lin-53* null allele *(bar19)* (Fig. S1E and Table S1). Interestingly, the short lifespan of *lin-53* mutants is not rescued upon overexpression of *lin-53* in muscles (*myo-3p::lin-53*), which, on the other hand, rescues the motility defect as shown earlier (Figs. 1F and 1H). In contrast, ubiquitous expression of recombinant LIN-53 using the *baf-1* promoter (*baf-1p::lin-53*; Fig. 1H) rescues the short lifespan in *lin-53* mutants confirming the functionality of heterologously expressed LIN-53 fusion proteins.

Our observations suggest that *lin-53* is required for healthy locomotion and a normal lifespan, raising the possibility that *lin-53* links lifespan regulation with healthspan maintenance.

### The muscle defect of *lin-53* mutants is phenocopied upon loss of the NuRD complex

LIN-53 is part of several different chromatin-regulating complexes including the CAF-1, NuRD, Sin3 and DRM complexes (Lu & Horvitz, 1998) (Loyola & Almouzni, 2004) (Eitoku et al., 2008). Hence, we wondered whether the observed phenotypes in *lin-53*-depleted animals are due to the altered function of a distinct complex. We generated an RNAi sub-library targeting all known LIN-53 interaction partners and tested whether depletion of any of the interaction partners phenocopies the observed muscle defects based on the *Pmyo-3::GFP* reporter (Fig. 2A). An impairment of muscle integrity became evident upon knock-down of genes encoding for NuRD complex members such as *lin-61*, *lin-40*, *dcp-66,* and, to a lesser degree, *let-418* and its paralog *chd-3* (Fig. 2A and 2B). This observation was further confirmed by immuno-staining of myosin (MHC) in mutant animals for NuRD-complex members (Fig. S2A), and motility assays (Fig. S2B) Overall, these findings suggested that the muscle phenotypes caused by depleting *lin-53* are due to affecting the NuRD complex. To test whether LIN-53 physically associates with the NuRD complex in muscles, we performed co-immunoprecipitation experiments coupled to mass spectrometry (IP-MS) using muscle-specific expression of FLAG-tagged LIN-53. Our muscle-specific IP-MS results revealed that LIN-53 interacts solely with chromatin-regulators that are part of the NuRD complex (Fig. S2C), strongly suggesting that LIN-53 associates with NuRD in muscles.

**Fig. 2:**
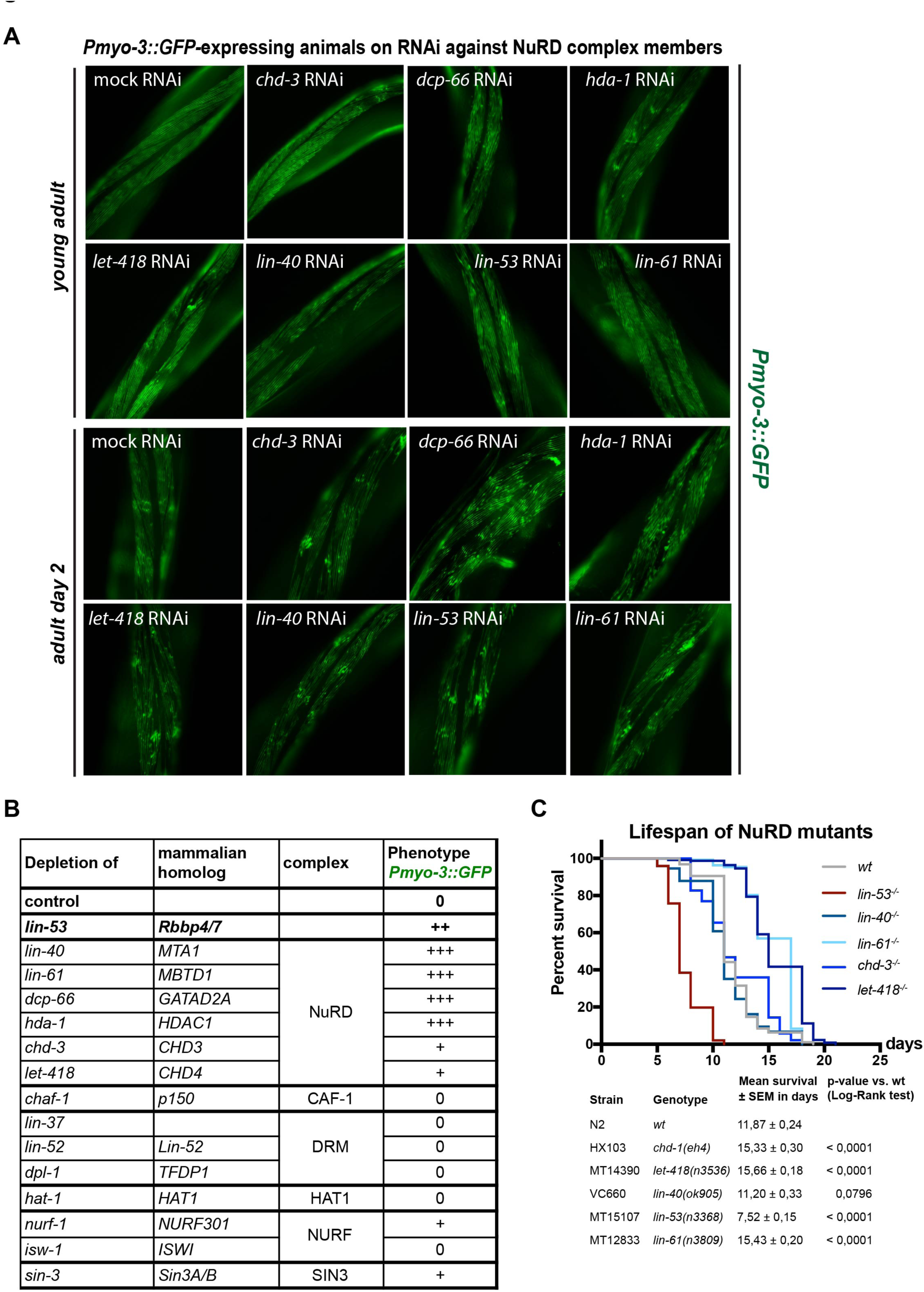
Loss of NuRD complex members phenocopy muscle defects of *lin-53* mutants. (A) Screening whether RNAi against genes encoding LIN-53-interacting phenocopy muscle integrity disruption as seen in *lin-53* mutants based on *Pmyo-3::GFP*. Representative fluorescent pictures of the *Pmyo-3::GFP* reporter are shown. (B) RNAi-depletion of NuRD complex members phenocopies *lin-53^−/-^* muscle phenotype. Disruption of the muscle-structure was scored: 0 = no effect, + = slight effect, ++ = medium defect, +++ = strong defect. (C) Depletion of NuRD complex members does not phenocopy the short lifespan of *lin-53* mutants. Survival analysis was carried out using Kaplan-Meier-estimator, p-value was calculated using Log-Rank Test. The experiment was done at least two times with at least 40 animals per repeat.

Next, we tested whether loss of the NuRD complex would also phenocopy the short lifespan of *lin-53* mutants. Surprisingly, depletion of NuRD members does not affect the lifespan of *C. elegans* but tends to rather increase lifespan as seen for *let-418* mutants (Fig. 2C). This observation is in agreement with a previous report showing that *let-418* mutants display an extended lifespan (De Vaux et al., 2013). Hence, in muscles, LIN-53 operates as part of the NuRD complex to maintain muscle integrity but does not seem to function through NuRD to ensure a normal lifespan of the animals.

### Short lifespan of *lin-53* mutants is phenocopied by *sin-3* mutants

The observation that LIN-53 is required for muscle maintenance but not lifespan regulation through the NuRD complex suggested that *lin-53* links healthspan with lifespan maintenance through different chromatin-regulating complexes. To identify through which complex LIN-53 maintains normal lifespan, we screened for a phenocopy of the short lifespan as seen in *lin-53* mutants, using available mutants of known LIN-53 interacting factors (Fig. 3). A phenocopy of the short lifespan as seen for *lin-53* mutants was only detectable in *sin-3* mutants (Fig. 3A), but not upon loss of any other known gene encoding a LIN-53-interacting protein (Figs. 3B - 3E). In *C. elegans*, the *sin-3* gene encodes the core subunit of the Sin3 chromatin-regulating complex, indicating that, in *lin-53* mutants, the integrity of the Sin3 complex might be affected, thereby causing the observed shortening of lifespan. This conclusion is further supported by the fact that lifespan is not further decreased upon depletion of *lin-53* in *sin-3* mutants arguing that both factors are involved in the same regulatory context (Fig. S3A). Hence, LIN-53 and SIN-3 cooperate to ensure normal lifespan in *C. elegans*.

**Fig. 3:**
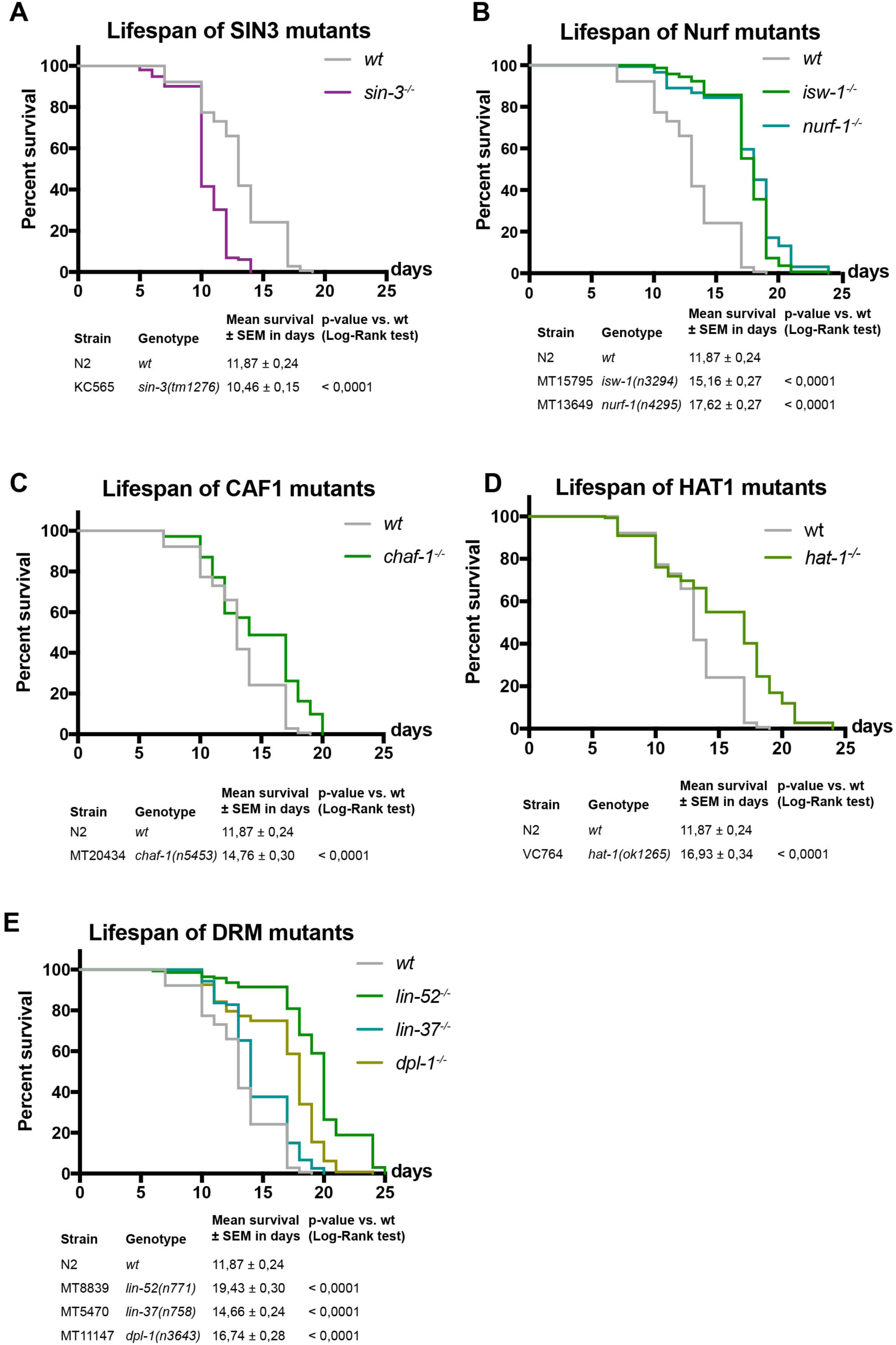
Lifespan of mutants for LIN-53-interactors. (A) Depletion of Sin3 complex member SIN-3 leads to shortened lifespan (violet line; mean lifespan 10,46 ± 0,15 days, p-value < 0,0001; compared to control; grey line; mean lifespan 13,66 ± 0,26 days. Survival analysis was carried out using Kaplan-Meier-estimator, p-value was calculated using Log-Rank Test. The experiment was done at least three times with at least 40 animals per repeat. (B - E) Lifespan analysis of different mutants from the DRM, Nurf, CAF1 and HAT1 complex. Survival analysis was carried out using Kaplan-Meier-estimator, p-value was calculated using Log-Rank Test. The experiment was done at least three times with at least 40 animals per repeat.

In summary, LIN-53 is interacting with the NuRD complex in muscles where its loss leads to a disruption of muscle integrity accompanied by locomotion defects. While affecting the NuRD complex does not lead to a short lifespan, loss of the Sin3 core subunit shortens lifespan, suggesting that LIN-53 regulates muscle homeostasis as part of the NuRD complex independently of lifespan regulation, which occurs through the Sin3 complex.

### Transcriptome of *lin-53* mutants shows mis-regulated metabolic genes

The shortened lifespan upon loss of LIN-53 suggested that specific molecular pathways might be affected in *lin-53* mutants. In order to examine this possibility, we performed whole transcriptome sequencing (RNA-Seq) and used both *lin-53* mutant backgrounds *n3668* (balanced) and *bar19* (CRISPR allele) (Fig. 4A). Our analysis revealed that 5.799 genes are differentially expressed in both *lin-53* mutant backgrounds when compared to the transcriptome of wild-type N2 animals (Figs. 4B and S4A). A number of muscle-related genes such as *hlh-1*, *unc-120*, *unc-52*, and *myo-3* are mis-regulated, which corresponds to the described motility defects in *lin-53* mutants (Fig. S4B). Interestingly, in both *lin-53* mutant backgrounds GO analysis (KEGG pathways) revealed a strong enrichment for differentially expressed genes that play a role in metabolic pathways (Figs. 4C and 4D). Since loss of SIN-3 phenocopies the short lifespan of *lin-53* mutants we also performed RNA-Seq analysis of the *sin-3(tm1276)* mutant background (Fig. 4E). Compared to *lin-53* mutants more than 50% of the differentially expressed genes in *sin-3* mutants overlap with those detected in both *lin-53* mutant backgrounds (Fig. 4E). Strikingly, GO analysis revealed a strong enrichment for genes that play a role in metabolic pathways also for *sin-3* mutants (Fig. 4F) suggesting that LIN-53 cooperates with SIN-3 in order to regulate metabolism. To elucidate whether LIN-53 might directly be involved in regulating the expression of ‘metabolic’ genes, we performed Chromatin Immunoprecipitation with subsequent sequencing (ChIP-Seq) using anti-LIN-53 antibody (Fig. 4G). The ChIP-Seq analysis revealed that the primary enriched pathway for genes which are bound by LIN-53 and become down-regulated upon loss of LIN-53, are implicated in metabolic pathways (Fig. 4G). This finding further corroborates the notion that LIN-53 is important for maintaining expression of genes that are important for metabolic processes. Since it is well established that metabolome alterations have a significant impact on aging (reviewed in (Peleg, Feller, Ladurner, & Imhof, 2016), (Finkel, 2015)), we propose that LIN-53 is required for normal lifespan of *C. elegans* because it is maintaining the expression of genes that ensure a wild-type metabolome.

**Fig. 4:**
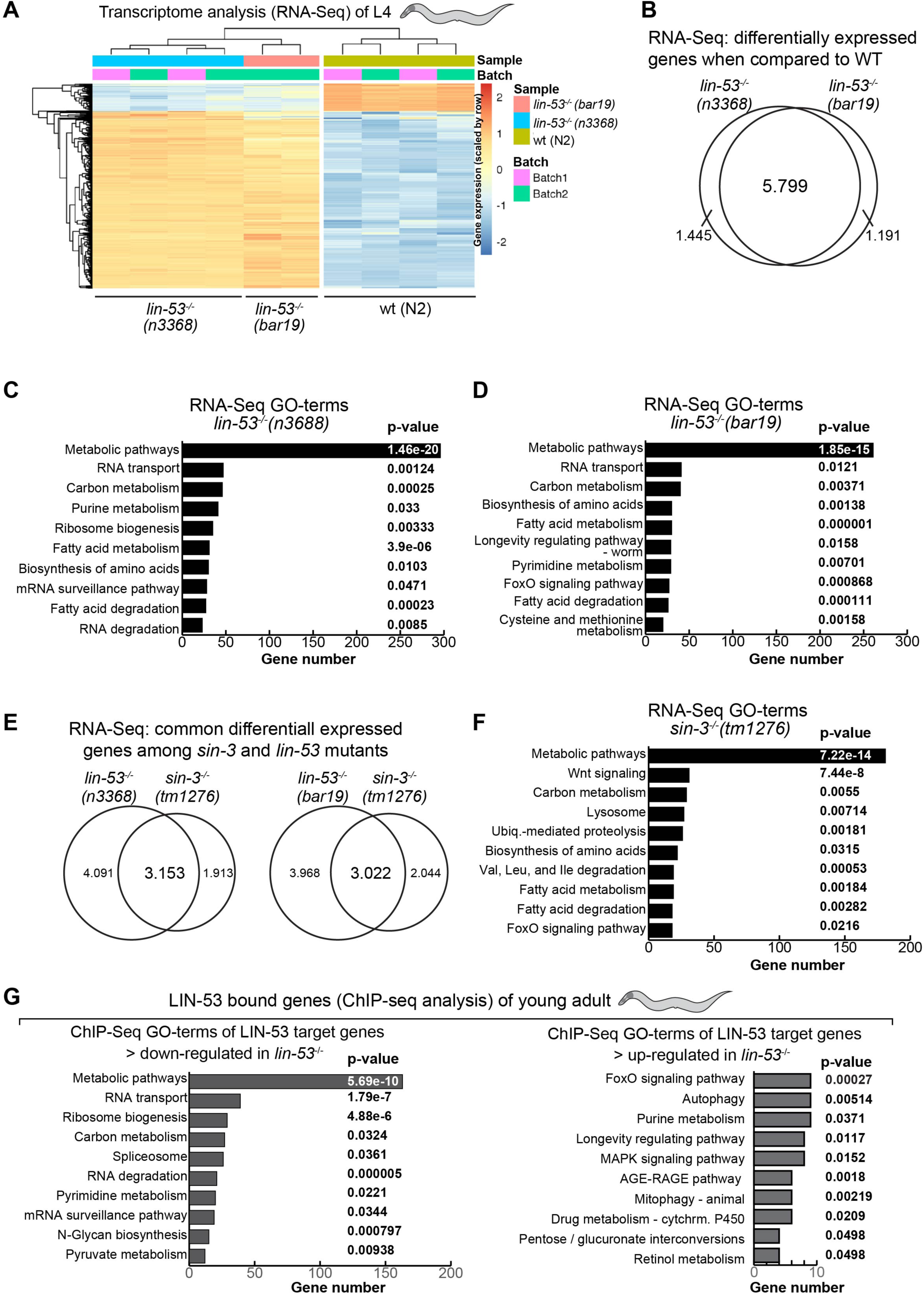
Differentially expressed genes in *lin-53* and *sin-3* mutants. (A) Heat-map of the normalized expression values (VST) of the Top 100 Genes with highest variance across samples in *lin-53(n3368)* and *lin-53(bar19)* mutants compared to control animals. (B) Venn diagram of differentially expressed genes in *lin-53(n3368)* and *lin-53(bar19)* mutants showing more than 5.000 overlapping genes. (C-D) Gene Ontology (GO) term analysis based on KEGG pathways of *lin-53* mutants compared to control animals using PANTHER. Mainly genes involved in metabolic processes are affected upon loss of *lin-53*. (E) Venn diagram of differentially expressed genes in *lin-53* and *sin-3* mutants. (G) GO-term analysis of ChIP-Seq results using PANTHER GO-Slim biological processes (KEGG pathways) for genes that are bound by LIN-53 and are either up regulated or down-regulated.

### Loss of *lin-53* leads to decreased levels of Trehalose

Next, we aimed to assess whether loss of LIN-53 leads to specific changes in the metabolome as suggested by the transcriptome and ChIP-Seq analyses. We examined the metabolome of *lin-53* and *sin-3* mutants at the young adult stage using Gas Chromatography coupled to Mass Spectrometry (GC-MS) and MS data analysis using Maui-SILVIA (Kuich, Hoffmann, & Kempa, 2014) (see methods) (Fig. 5A). Wild-type animals were used as a control, as well as *let-418* mutants (NuRD complex). Since *let-418* mutant animals do not have a shortened lifespan (Fig. 2C), metabolites that change in *lin-53* and *sin-3* mutant animals, but not in *let-418* mutants, are likely to be implicated in the short lifespan phenotype upon loss of *lin-53* or *sin-3*. The unique metabolite, which showed such a pattern was the glucose disaccharide Trehalose. Trehalose levels are decreased in *lin-53* and *sin-3* mutants but not in *let-418* mutants (Fig. 5A). Interestingly, it has previously been shown that decreased Trehalose levels lead to a shortened lifespan in *C. elegans* (Y. Honda et al., 2010), (Seo et al., 2018) suggesting that reduced Trehalose levels in *lin-53* and *sin-3* mutants may cause the observed short lifespan of these animals. Impaired maintenance of Trehalose levels is also reflected by the fact that reporter expression for the Trehalose 6-Phosphate Synthase-encoding genes *tps-1* and *tps-2* (Y. Honda et al., 2010), which are essential for Trehalose synthesis, are reduced upon knock-down of *lin-53* (Fig. 5B, and Fig. S5A). Analysis by qRT-PCR (Fig. 5B) confirmed a down regulation of *tps-1* in short-lived *lin-53* and *sin-3* mutants, but not in long-lived *lin-40* and *let-418* mutants.

**Fig. 5:**
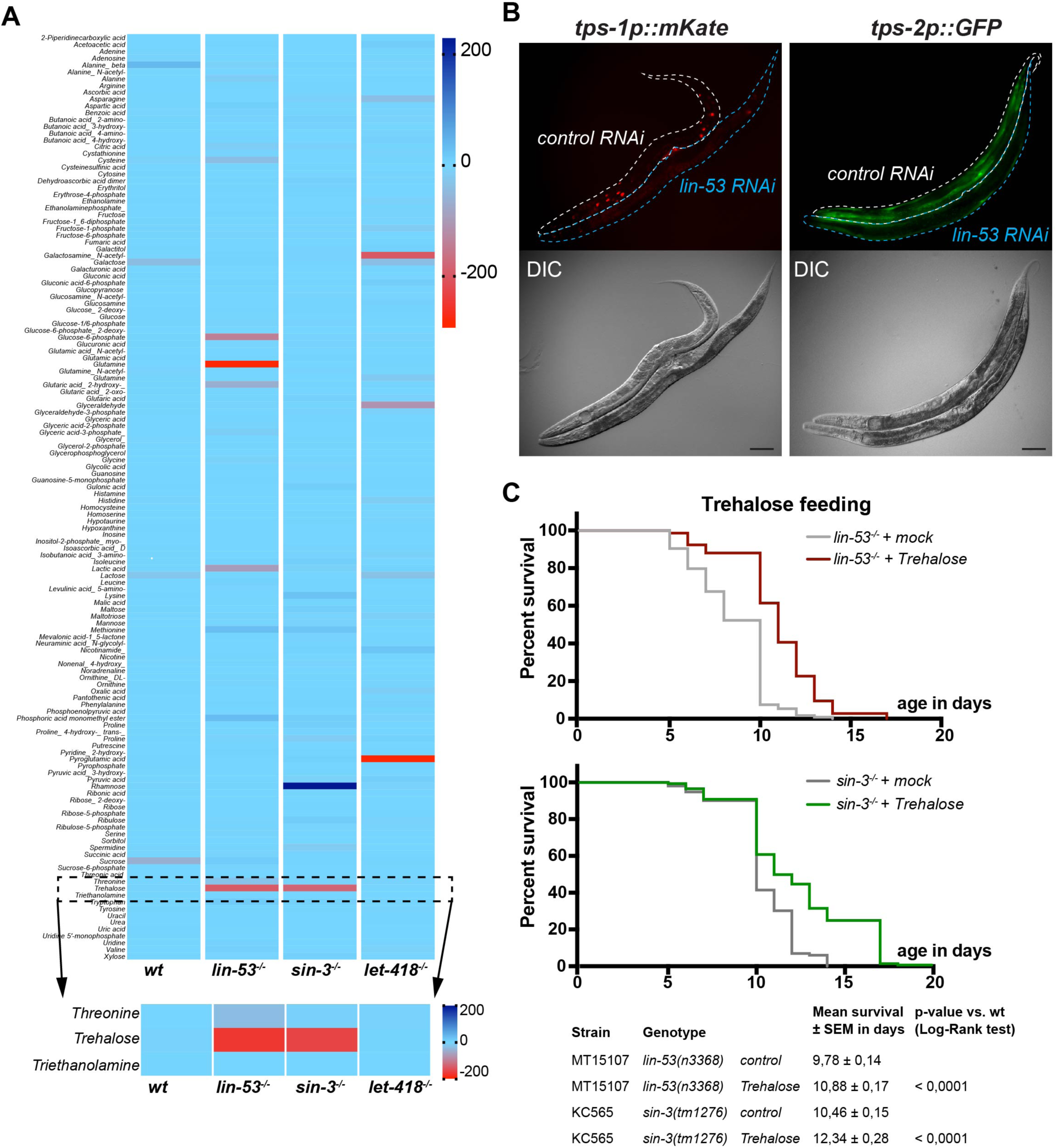
Metabolome analysis reveals decreased Trehalose biosynthesis in mutants. (A) Metabolome analysis of wt, *lin-53^−/-^*, *sin-3^−/-^,* and *let-418^−/-^* mutants. 215 different metabolites were detected using GC-MS. The data was z-transformed and plotted as a heat-map. Trehalose is decreased in *lin-53* and *sin-3* mutants, but not changed in wt and long-lived *let-418* mutants. 2 biological repeats were analyzed. (B) Depletion of *lin-53* depletion leads to decreased expression of *tps-1p::mKate* and *tps-2p:.GFP.* Control and *lin-53* RNAi-treated worms were mounted next to each other allowing direct comparison. Upon *lin-53* RNAi animals show a decrease in expression of both reporters. (C) Short lifespan of *lin-53* and *sin-3* mutants is partially rescued after feeding with Trehalose (mean lifespan of *lin-53* mutants on trehalose 10,98 ± 0,13 days; *lin-53* mutants 8,58 ± 0,13 days, p-value < 0,0001; mean lifespan of *sin-3* mutants on Trehalose 10,46 ± 0,15 days; *sin-3* mutants 10,46 ± 0,15 days, p-value < 0,0001). The experiments were carried out three times with at least 40 animals scored per repeat. Survival analysis was done using Kaplan-Meier-estimator, p-value was calculated using Log-Rank Test.

To provide further evidence that Trehalose reduction contributes to shortening the lifespan in *lin-53* and *sin-3* mutants, we tested whether feeding of Trehalose would alleviate the aging phenotype (Fig. 5C). Replenishing *lin-53* and *sin-3* mutants with Trehalose by feeding resulted in extended lifespans compared to unfed mutants (Fig. 5C) indicating that reduced levels of Trehalose play a role in shortening the lifespan upon loss of *lin-53*.

### Loss of LIN-53 affects the Insulin signaling (IIS) pathway

Recently, it has been demonstrated that the Insulin/IGF1 signaling (IIS) pathway is important to promote the benefits of Trehalose in the context of lifespan maintenance (Seo et al., 2018). It is important to note in this context that animals carrying loss of function alleles of the *daf-2* gene, which encodes the IIS receptor, were identified as one of the first mutants with significantly extended lifespans (reviewed in (Kenyon, 2011)) and it has been reported that Tehalose synthesis is up-regulated in *daf-2* mutants (Hibshman et al., 2017; Y. Honda et al., 2010).

In order to test whether LIN-53 is implicated in regulating Trehalose levels via the IIS pathway we first assessed the lifespan of *lin-53(n3368)*; *daf-2(e1370)* double mutants (Fig. 6A). While the *lin-53(n3368)*; *daf-2(e1370)* double mutants live longer than *lin-53* mutants alone, which is comparable to the lifespan of wild-type animals, they live significantly shorter than *daf-2* mutants alone suggesting a requirement for LIN-53 in IIS pathway-mediated lifespan extension (Fig. 6A). Similar results were obtained when comparing lifespans of the double *sin-3(tm1276); daf-2(e1370)* animals with the respective single mutants (Fig. 6B). These observations suggested that the previously reported increase of Trehalose levels in *daf-2* mutants (Y. Honda et al., 2010) might compensate for the diminished Trehalose levels upon loss of LIN-53 and SIN-3. To test this assumption, we analyzed the metabolome of *lin-53(n3368)*; *daf-2(e1370)* as well as *sin-3(tm1276); daf-2(e1370),* and found that Trehalose levels in both double mutants are similar to that of wild-type animals (Fig. 6C). We therefore concluded that *daf-2* mutants suppress the short lifespan of *lin-53* and *sin-3* mutants by counteracting the Trehalose deprivation upon loss of LIN-53 or SIN-3.

**Fig. 6:**
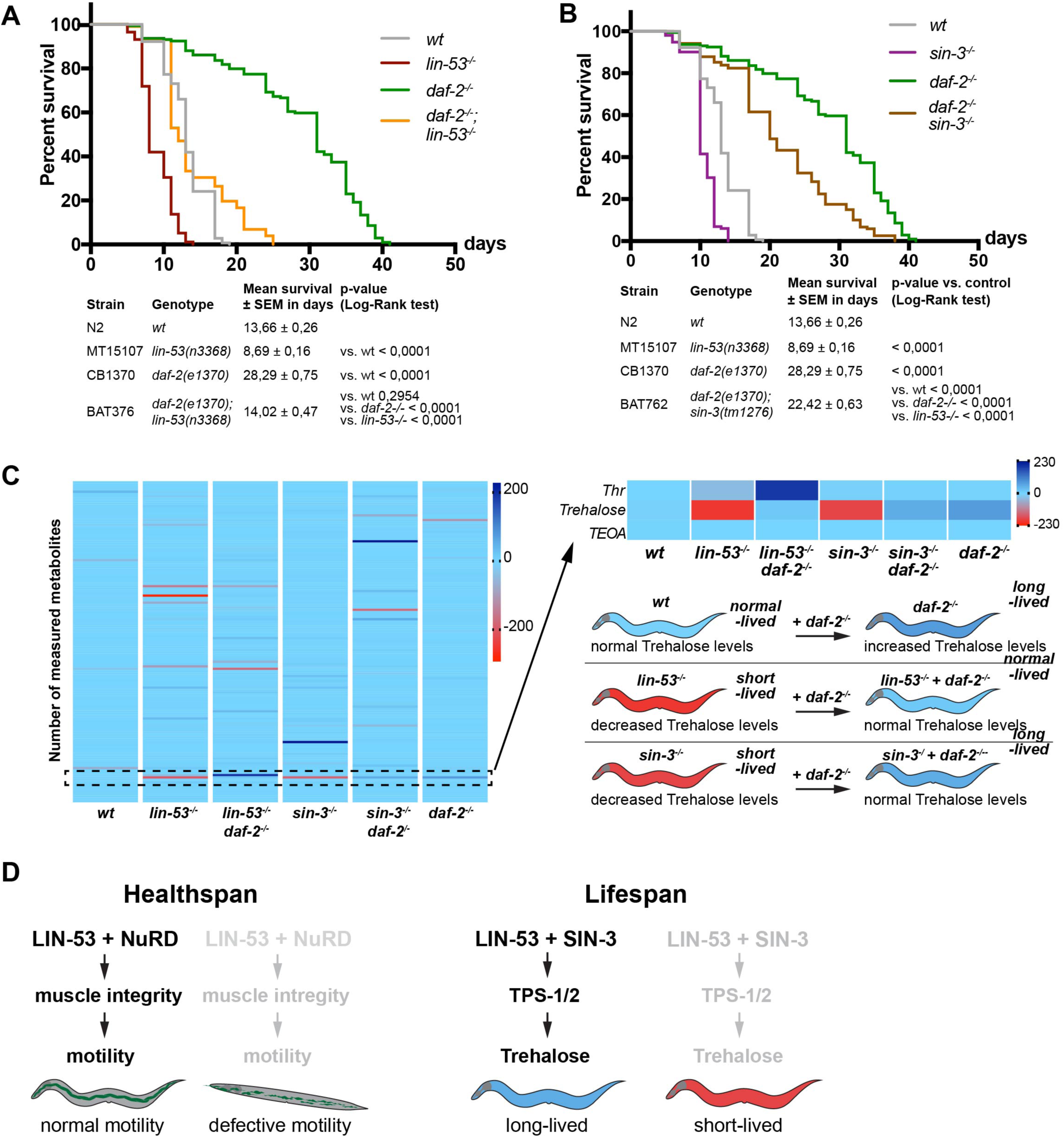
The Insulin/IGF1 receptor mutant *daf-2(e1370)* restores Trehalose levels. (A) The short lifespan of *lin-53* mutants is partially rescued by *daf-2*. wt animals (grey line; mean lifespan 13,66 ± 0,26 days), *lin-53* mutants (red line; mean lifespan 8,69 ± 0,16 days), *daf-2* mutants (green line; mean lifespan 28,29 ± 0,75 days), *daf-2; lin-53* double mutants (orange line; mean lifespan 14,02 ± 0,47 days). Triplicate experiments were carried out with 40 animals per repeat. Survival analysis was carried using Kaplan-Meier-estimator, p-value was calculated using Log-Rank Test. (B) As seen for *lin-53* mutants, short lifespan of *sin-3* mutants is suppressed by the *daf-2(e1370)* mutation. wt animals (grey line; mean lifespan 13,66 ± 0,26 days), *sin-3* mutants (violet line; mean lifespan 10,46 ± 0,15 days), *daf-2* mutants (green line; mean lifespan 28,29 ± 0,75 days), *daf-2 sin-3* double mutants (brown line; mean lifespan 22,42 ± 0,63 days). Triplicate experiments were carried out with 40 animals per repeat. Survival analysis was carried using Kaplan-Meier-estimator, p-value was calculated using Log-Rank Test. (C) Metabolome analysis of wt, *lin-53^−/-^*, *daf-2^−/-^, lin-53^−/-^; daf-2^−/^, sin-3^−/-^, sin-3^−/-^* and *daf-2^−/-^* mutants. Trehalose levels are increases in long-lived *daf-2 (e1370)* mutants. The GC-MS data was z-transformed and plotted in a heat-map. Trehalose is increased in *daf-2* and restored in *daf-2; lin-53* double mutants as well as *sin-3; daf-2* double mutants. 2 biological repeats were analyzed. (D) Model summarizing the findings of LIN-53 implication if lifespan and healthspan regulation.

Overall, our findings suggest that LIN-53 maintains sufficient Trehalose levels to ensure a normal lifespan in conjunction with SIN-3 and this maintenance has interplay with the IIS pathway in *C. elegans*. Moreover, the muscle defects and short lifespan in *lin-53* mutants can be unlinked: LIN-53 is interacting with the NuRD complex to maintain muscles and proper motility but ensures normal lifespan via the Sin3 complex (Fig. 6D). Loss of LIN-53 or SIN-3 leads to diminished levels of the disaccharide Trehalose, which is required for a normal lifespan (Y. Honda et al., 2010),(Seo et al., 2018). These findings suggest that the histone chaperone LIN-53 is a critical chromatin regulator linking the epigenetic regulation of healthspan and lifespan.

## Discussion

Recent studies revealed epigenetic factors as an emerging group of aging regulators that control gene expression at the level of chromatin (reviewed in (Brunet & Rando, 2017)). For instance, the ASH-2 trithorax complex regulates aging in *C. elegans* by catalyzing histone H3 methylation at Lysine residue 4 (K4) (Greer et al., 2010) and loss of epigenetic regulation in mouse hematopoietic stem cells accelerates aging (Chambers et al., 2007). In the context of aging regulation, one important aspect is whether healthspan and lifespan are intimately linked. Meaning, should we expect that animals or humans with longer lifespans would also be healthy for a longer time? Interestingly, a recent study by the research group of Heidi Tissenbaum suggests, that lifespan and healthspan can be unlinked in *C. elegans* (Bansal, Zhu, Yen, & Tissenbaum, 2015). For instance, the authors showed that in many cases, when lifespan is extended, there is an increase in the time for which animals live in a frail state (Bansal et al., 2015). As many aging-regulating pathways are evolutionarily conserved, an unlinking of lifespan and healthspan is conceivable also in higher organisms. Our findings described in this study corroborate this notion as the histone chaperone LIN-53 is highly conserved in metazoan species (known as RBBP4/7 and CAF-1p48 in mammals) (Harrison et al., 2006). We found that overexpression of the chromatin regulator LIN-53 specifically in muscles of *C. elegans*, rescued the *lin-53* null-mutant phenotype with regard to the muscle and motility defects. Strikingly, high levels of LIN-53 in muscles are beneficial because motility remained in a healthy wild type-like state even in aged animals. However, while muscle-specific LIN-53 overexpression reconstituted muscle health in *lin-53* mutants, the lifespan of these animals remained short, which suggested that the effects of LIN-53 on muscle health and lifespan are separable. Our finding that LIN-53 associates with the NuRD complex in muscles in order to maintain muscle integrity, while its role in lifespan homeostasis is mediated via the Sin3 complex, confirmed this initial assumption. RNA-Seq analysis of mutant *lin-53* animals further indicates a global mis-regulation of muscle genes such as the myosin heavy chain-encoding genes *unc-54* and *myo-3* as well as muscle transcription factors including HLH-1 and HND-1, which could explain the observed muscle and motility phenotypes upon loss of *lin-53* and NuRD components. Notably, mutants for the NuRD subunit LET-418 have an increased lifespan, but still show a compromised movement suggesting that loss of the NuRD complex only affects muscle integrity. While the exact molecular mechanism by which LIN-53 regulates muscle homeostasis remains to be determined, our findings provide an important initial framework for elucidating LIN-53’s roles in muscles via the NuRD complex.

With respect to the aging phenotype of *lin-53* mutants we found that the shortened lifespan is caused by loss of the Sin3 complex. Animals deleted for the *sin-3* gene phenocopy not only the short lifespan of *lin-53* mutants but also show similar patterns of gene expression changes when compared to wild-type animals. The fact that loss of either *lin-53* or *sin-3* primarily affects mainly the expression of genes related to metabolic processes prompted us to assess changes in the metabolome in these mutants. Strikingly, our analysis showed that Trehalose levels are diminished in both *lin-53* and *sin-3* mutants, thereby revealing a possible common factor with regard to impacting lifespan regulation. It is known that decreased Trehalose levels result in a shorter lifespan in *C. elegans*, as, described earlier (Y. Honda et al., 2010). Our finding that Trehalose levels are reconstituted when we combined either *lin-53* or *sin-3* mutants with the *daf-2* mutant background is, therefore, in agreement with previous studies showing that the insulin/IGF-signaling (IIS) pathway controls Trehalose levels and that loss of DAF-2 results in increased Trehalose levels (Y. Honda et al., 2010).

A conserved role for LIN-53 in aging regulation is conceivable because its human homologs RBBP4 and RBBP7 have been implicated in Hutchinson–Gilford Progeria Syndrome (HGPS), which leads to premature aging (Pegoraro et al., 2009). HGPS belongs to laminopathic disorders caused by mutations in genes encoding for lamin A/C or for other nuclear lamina proteins such as Emerin (Zaremba-Czogalla, Dubińska-Magiera, & Rzepecki, 2010). In primary dermal fibroblasts of HGPS patients, RBBP4/7 levels are significantly reduced, which is also the case in fibroblasts from aged human beings (Pegoraro et al., 2009). However, in their study Pegoraro et al. proposed that the premature aging disorder is caused by the loss of functional NuRD complexes due to reduced levels of RBBP4/7 (Pegoraro et al., 2009). While we do not see premature aging upon depletion of specific NuRD subunits such as LIN-40, LIN-61, or LET-418 in *C. elegans,* we identified the Sin3 complex to be relevant for the aging phenotype upon loss of LIN-53. We speculate that the Sin3 complex might also play a role during aging regulation in other species as it has also been shown by a previous study in *Drosophila* that knock-down of the Sin3A gene causes a shortened lifespan (Barnes et al., 2014).

In humans LIN-53 homologs might link aging regulation with healthspan as we see it in the *C. elegans*. Laminopathies such as the Emery–Dreifuss Muscular Dystrophy (EDMD) diminish muscle maintenance and motility in patients (Zaremba-Czogalla et al., 2010). Laminopathies in general might affect RBBP4/7 levels as shown in the laminopathic disorder HGPS thereby recapitulating the phenotypes of *lin-53* mutant worms. Hence, reduced levels of LIN-53 and its homologs RBBP4/7 might lead to premature aging and impaired maintenance of muscle integrity in *C. elegans* as well as humans. Interestingly, it has recently been revealed that the decline of RBBP4 in humans causes cognitive aging and memory loss (Pavlopoulos et al., 2013). Hence, it is conceivable that the role of LIN-53 as a link of lifespan regulation with healthspan maintenance might be conserved also in higher organisms, which can have important implications for human health during aging.

## Experimental Procedures

List of strains used in this study.

| Strain name | Genotype | Reference |
| --- | --- | --- |
| N2 | *wt* | CGC |
| MT15107 | *lin-53(n3368)* | CGC |
| CB1370 | *daf-2(e1370)* | CGC |
| BAT376 | *daf-2(e1370); lin-53(n3368)* | this study |
| BAT1883 | *lin-53(n3368); barEx974 [baf-1p::GFP::lin-53::2xFLAG]* | this study |
| BAT729 | *lin-53(n3368); barIS87 [myo-3p::lin-53::2xFLAG]* | this study |
| HX103 | *chd-3(eh4)* | CGC |
| VC924 | *dcp-66(gk370)* | CGC |
| MT14390 | *let-418(n3536)* | CGC |
| VC660 | *lin-40(ok905)* | CGC |
| MT12833 | *lin-61(n3809)* | CGC |
| MT15795 | *isw-1(n3294)* | CGC |
| MT13649 | *nurf-1(n4295)* | CGC |
| MT8839 | *lin-52(n771)* | CGC |
| MT5470 | *lin-37(n758)* | CGC |
| MT11147 | *dpl-1(n3643)* | CGC |
| MT20434 | *chaf-1(n5453)* | CGC |
| KC565 | *sin-3(tm1276)* | CGC |
| VC764 | *hat-1(ok1265)* | CGC |
| BAT762 | *daf-2(e1370); sin-3(tm1276)* | this study |
| BAT1368 | *myo-3p::lin-53_IR::2xNLS::tagRFP* | this study |
| BAT1265 | *myo-3p::lin-53::GFP::2xFLAG::SL2::tagRFP line 2* | this study |
| BAT1982 | *tps-1p::mKATE::H2B* | this study |
| BC18476 | *tps-2p::GFP* | CGC |

List of RNAi clones used in this study.

| Gene name | derived from |
| --- | --- |
| empty vector | Addgene |
| *lin-53* | Tursun *et al.*, 2012 |
| *chd-3* | Chromatin 2.0 library from (Hajduskova *et al.*, 2018) |
| *dcp-66* |  |
| *hda-1* |  |
| *let-418* |  |
| *lin-40* |  |
| *lin-61* |  |

List of primers for qPCR

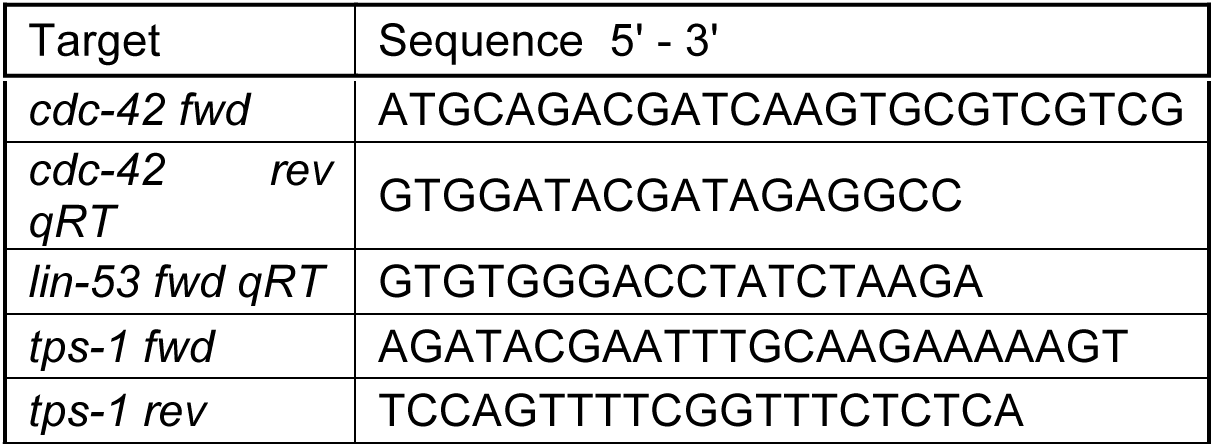

### Lifespan and lococmotion assay

For age synchronization, eggs were put on a plate, which was scored as day 0. Worms were grown until L4 stage at 15°C and then transferred to plates containing 5-Fluoro-2′-deoxyuridine (FUDR; 10 worms per plate) and further grown at 25°C. The animals were scored daily for survival and the locomotion was classified (categories A – C) (Herndon et al., 2002). Animals that did not show pharyngeal pumping or respond to podding were scored as dead. For data analysis of the survival OASIS was used (Han et al., 2016).

### Thrashing assay

Age synchronized worms were put in a drop of 10 µl M9 buffer and a video was made using the DinoXcope software in combination with a Dino-Lite camera. The video was taken with a resolution of 640×480 with a frame rate of 24.00 fps for 30 seconds at normal quality. The calculation of the body bends was carried out using the ImageJ Plugin wrMTrck (Jesper S. Pedersen) with the according settings.

### Immunostaining and antibodies

Staining was carried out as previously described (Seelk et al., 2016). In brief, worms were freeze-cracked after resuspension in 0,025 % glutaraldehyde between two frost-resistant glass slides on dry ice. The animals were fixed using Acetone/Methanol for 5 min each and washed off into PBS. Afterwards the sample was blocked in 0,25 % Triton + 0,2 % Gelatine in PBS and stained. Primary antibodies were diluted in PBS with 0,25 % Triton + 0,1 % Gelatine and the fixed worms were incubated overnight at 4°C. After PBS washes secondary antibody was added for 3 hrs. Worms were mounted with DAPI-containing mounting medium (Dianova, #CR-38448) on glass slides after further washing steps. The primary antibodies used were anti-MHC (1:300; DHSB #5-6-s) and anti-LIN-53 (1:800, Pineda). As Secondary antibodies Alexa Fluor dyes were applied at 1:1000 dilution.For phalloidin staining the worms were harvested with PBS and fixed with 4% formaldehyde in PBS. For freeze-cracking worms were frozen in liquid nitrogen followed by thawing at 4°C for three times. The samples was incubated for 30 min followed by three times washing with PBST for 10 min. Phalloidin-rhodamine in PBST was added and incubated for 30 min on room temperature. After a last washing for three 10 min in PBST the worms were mounted on slide using mounted with DAPI-containing mounting medium (Dianova, #CR-38448). Microscopy was done using the Zeiss Axio Imager 2 fluorescent microscope.

### RNA interference

RNA inference was usually carried out as P0, meaning that eggs were put on RNAi plates and the same generation was scored. Worms were bleached and eggs were put on RNAi plates seeded with bacteria expressing dsRNA or carrying an empty RNAi vector and grown at 15°C until they reached L4 stage. If animals were used for a lifespan experiment, 10 animals were put on one RNAi plate containing FuDR and cultured further at 25°C. For monitoring of the muscle structure worms were grown at 15°C until they reached the stage of interest and analyzed by fluorescent microscopy. For construction of the *lin-53* interaction partner sublibrary, candidate genes interacting with LIN-53 were chosen based on a literature search (www.pubmed.com). The library was generated by compiling the clones from the chromatin RNAi sublibrary generated in Hajdoskova et al, 2018. The list of RNAi clones used can be found in table 0.

### Generation of a *lin-53* hairpin construct

For generation of a *lin-53* short hairpin construct (shRNA), the desired fragment was amplified using specific primers to introduce two different restriction sites at both ends of the cDNA of *lin-53* (Tavernarakis, Wang, Dorovkov, Ryazanov, & Driscoll, 2000). The restriction site is used as an inversion point to ligate two pieces of *lin-53* together to form an inverted repeat and clone into a plasmid carrying a muscle-specific promotor to enable expression of the *lin-53* shRNA in muscles.

### Co-Immunoprecipitation with subsequent Mass Spectrometry (IP-MS)

Worms were synchronized by bleaching and grown on 5 – 10 15 cm plates until L4 stage. Worms were washed off the plates using M9 buffer and freeze-cracked by adding the worm suspension dropwise to liquid nitrogen, pulverized using a hammer and a bio-pulverizer for 20 – 30 times and afterwards ground to a fine powder using a mortar. The powder was thawed and dissolved in 1,5 vol lysis buffer (50 mM HEPES-KOH (pH 7,6); 1 mM EDTA; 0,25 M LiCl; 1% sodium deoxycholate; 0,5 % NP-40; 100 mM NaCl; 10% Glycerol + 1 tablet of Complete in 10 ml of buffer). In order to shear DNA, the sample is sonicated using Bioruptor^®^ device for six times with 30 sec ON and 30 sec OFF on high settings. The lysate is cleared by spinning and µMACS-DKYDDDK (Milteny Biotec) are added and incubated for 30 min on ice, before the mixture is applied on a magnetic M column. After washing three times with lysis buffer the samples are eluted using 8 M guanidiniumhypochlorid pre-heated to 80°C for mass spectrometry (MS). Sample preparation for MS was done as described previously (Hajduskova et al., 2018). Raw MS data was analyzed using MaxQuant Software (Cox & Mann, 2008).

### RNA extraction

Whole transcriptome sequencing was carried out as previously described (Kolundzic et al., 2018). In brief, RNA was extracted from control, *lin-53(n3368)*, *lin-53(bar19)* and *sin-3(tm1276)* animals using TRIzol (Life Technologies) and guanidinium thiocyanate-phenol-chloroform extraction. Adding chloroform to the TRIzol sample leads to a phase separation with the aqueous phase containing the RNA, an interphase and an organic phase containing DNa and proteins. The RNA was further purified from the aqueous phase using isopropanol.

### qRT-PCR

To analyze gene expression, RNA was first reverse transcribed using GoScript Reverse transcriptase (Promega) using oligo(dT) and random hexamer primers. The qPCR was carried out using Maxima SYBR Green/ROX qPCR Master Mix (2x) according to the manufacturer’s instructions. The measurement was done using the CFX96 Touch Real-Time PCR Detection System from BioRad. *cdc-42* was used as a reference gene and relative expression was calculated using the Livak method (Schmittgen & Livak, 2008).

### Whole transcriptome sequencing

Library preparation for RNA-Sequencing was carried out using TruSeq RNA Library Prep Kit v2 (Illumina) ccording to the manufacturer’s instructions. Libraries were sequenced using paired eind sequencing length of a 100 nucleotides on a HiSeq4000 machine (Illumina).

### Analysis of RNA-seq data

The RNA-seq sequencing datasets were processed using the PiGx-RNAseq (Wurmus et al., 2018) pipeline (version 0.0.4), in which the quality of the raw fastq reads were improved using Trim Galore (https://www.bioinformatics.babraham.ac.uk/projects/trim_galore/.), gene-level expression was quantified using Salmon (Patro, Duggal, Love, Irizarry, & Kingsford, 2017) based on worm transcript annotations from the Ensembl database (version 89). The raw read counts were further processed using RUVs function of the RUVseq R package (Risso, Ngai, Speed, & Dudoit, 2014) to remove unwanted variation from the expression data. Covariates discovered using RUVseq was integrated with DESeq2 (Love, Huber, & Anders, 2014) to test for differential expression using a lfcThreshold of 0.5 and false discovery rate of 0.05. The GO term enrichment is calculated using the gProfileR package. The p-values are corrected for multiple testing. The default multiple testing correction method (“analytical”) was used when running the gProfileR’s main enrichment function ‘gprofiler’.

### ChIP-Seq

In brief, in M9 arrested L1 worms were grown on OP50 plates for 40h to L4/YA stage at room temperature. Animals were washed three times with M9 and fixed with 2% formaldehyde for 30 minutes followed by quenching with 0.125M glycine for 5 minutes. The samples were rinsed twice with PBS, and 200-300 ul of pellets were snap-frozen in liquid nitrogen and kept at −80°C. The pellets were washed once with 0.5 ml PBS + PMSF and resuspended in 1 ml FA Buffer (50mM HEPES/KOH pH 7.5, 1mM EDTA, 1% Triton X-100, 0.1 sodium deoxycholate, 150mM NaCl)+0.1% sarkosyl+protease inhibitor (Calbiochem) and then dounce-homogenized on ice with 30 strokes. The samples were sonicated with Biorupter with the setting of high power, 4°C, and 15 cycles of 30 sec on 30 sec off. Soluble chromatin was isolated by centrifugating for 15 min at max speed and 4°C. The cellular debris was resuspended in 0.5 FA Buffer+0.1% sarkosyl+protease inhibitor and sonicated again 15 cycles with the same setting. Isolated soluble chromatin were combined. The immunoprecipitation of LIN-53 protein was performed overnight in a total volume of 600 µ l with 10 µl of PA58 (polyclonal peptide AB; rabbit, Pineda), while 5% of samples were taken as input. Immunocomplexes with collected with Protein A-Sepharose beads (Sigma). The beads were washed with 1 ml of following buffers: twice with FA Buffer for 5 min, FA-1M NaCl for 5 min, FA-0.5M NaCl for 10 min, TEL Buffer (0,25M LiCL, 1% NP-40, 1% Sodium deoxycholate, 1mM EDTA, 10mM Tris-HCl pH 8.0) for 10 min, and twice with TE Buffer (pH8.0). DNA-protein complexes were eluted in 250 ul ChIP elution buffer (1%SDS, 250mM NaCl, 10 mM Tris pH8.0, 1mM EDTA) at 65°C for 30 min by shaking at 1400 rpm. The Inputs were treated for approx. 3h with 20µg of RNAse A (Invitrogen). The samples and inputs were treated with 10µg of Proteinase K for 1h, and reverse cross-linked overnight at 65°C. DNA was purified with Qiagen MinElute PCR purification Kit. Sequencing library preparation was carried out using NEXTflex qRNA-seq Kit v2 Set A kit according to manufacturer’s instructions. Libraries were sequenced with Hiseq 4000 mit 2×75 bp.

### ChIP-seq sequencing data processing

The ChIP-seq sequencing datasets were processed using the PiGx-ChIPseq (Wurmus et al., 2018) pipeline (version 0.0.16), in which the quality of the raw fastq reads were improved using Trim Galore (https://www.bioinformatics.babraham.ac.uk/projects/trim_galore/.), processed reads were aligned to the DNA sequence assembly WBcel235 (Ensembl version 89) using Bowtie2 (Langmead & Salzberg, 2012), the peaks were called using MACS2 (Zhang et al., 2008), and the IDR (irreproducible discovery rate) peaks were called using the IDR software (Li, Qunhua Li, Brown, Huang, & Bickel, 2011).

### Data availability

To see a detailed description of the downstream analysis of RNA-seq and ChIP-seq data, see the github repository: https://github.com/BIMSBbioinfo/collab_seelk_tursun_lin53_paper

### Metabolome analysis

Wild type, *daf-2(e1370)*, *lin-53(n3368)*, *sin-3(tm1276)*, *daf-2(e1370); lin-53(n3368)* double mutants were analyzed at L4 stage. For sample-collection worms were synchronized by bleaching and transferred to NGM-plates containing OP50 as food source. Worms were grown at 15°C until L3/4 stage and then shifted to 25°C until young adult stage. Worms were harvested in M9 medium and adjusted to approx. 40 mg of worms per sample. The sample extraction is performed by using methanol::chloroform::water (5:2:1, MCW; 1 ml per 50 mg sample). To lyse worms and immediately bring the metabolites in solution 500 µl ice-cold MCW (with cinnamic acid) is added to the frozen worm sample. The sample was transferred to a new tube containing Silica beads and lysed using mechanical force with a tissue lyser at 6.500 ^m^/_s_, 2×20 sec ON, 5 sec OFF for three times. To further solubilize, the lysate was sonicated for 10 min in an ultrasound bath. As much supernatant as possible was taken off the beads and the left over MCW (x – 500 µl) was added, the sample was vortexed and shortly incubated on dry ice. After shaking for 15 min, 1.400 rpm at 4°C 0,5 vol of water were added for phase separation. The sample was vortexed and shaken for 15 min, 1.400 rpm at 4°C. Vortexing was repeated and the sample was spun at max. speed, 4°C for 10 min to ensure phase separation. The polar phase was taken off and further prepared for measurement. The polar phase was dried overnight in a speed vac followed by derivatization. For this, first 10 µl of 40 ^mg^/_ml_ methoxyamine hydrochloride (MeOx) solution in pyridine is added and incubated for 90 min at 30°C with shaking. Afterwards 30 µl N-Methyl-N-(trimethylsilyl) trifluoroacetamide (MSTFA) is added to the sample. Per 1.000 µl of MSTFA 10 µl of a standard retention index mixture of different decanes (C17 mix) is dissolved in MSTFA. Everything is incubated at 30°C for 1 h with shaking. After spinning down for 10 min at full speed the samples are put into glass vials for the GC-MS. The data analysis was carried out using Maui-SILVIA (Kuich et al., 2014).

## Acknowledgement

We thank ALina El-Khalili and Sergej Herzog and Tim Wolfram for technical assistance. Also, we are grateful for the support by Matthias Selbach and Stefan Kempa for proteomics and metabolomics applications. We thank the CGC, supported by the NIH, for providing strains, Ena Kolundic, Martina Hajduskova and Anna Reid for discussion and comments on the manuscript. All procedures conducted in this study were approved by the Berlin State Department for Health and Social (LaGeSo).

## Author contribution

SM and BT designed the study and interpreted the results. SM, AK, BV, SB, and MH conducted experiments and analyzed data. SM and BT wrote the manuscript. BU and AA performed bioinformatic analyses. BT acquired funding for the project from the ERC. All authors assisted in editing the manuscript.

## Conflict of interest

The authors declare that they have no competing interests.

## Funding

This work was partly sponsored by the ERC-StG-2014-637530 and ERC CIG PCIG12-GA-2012-333922 and is supported by the Max Delbrueck Center for Molecular Medicine in the Helmholtz Association.

## Supporting Information

**Fig. S 1.**
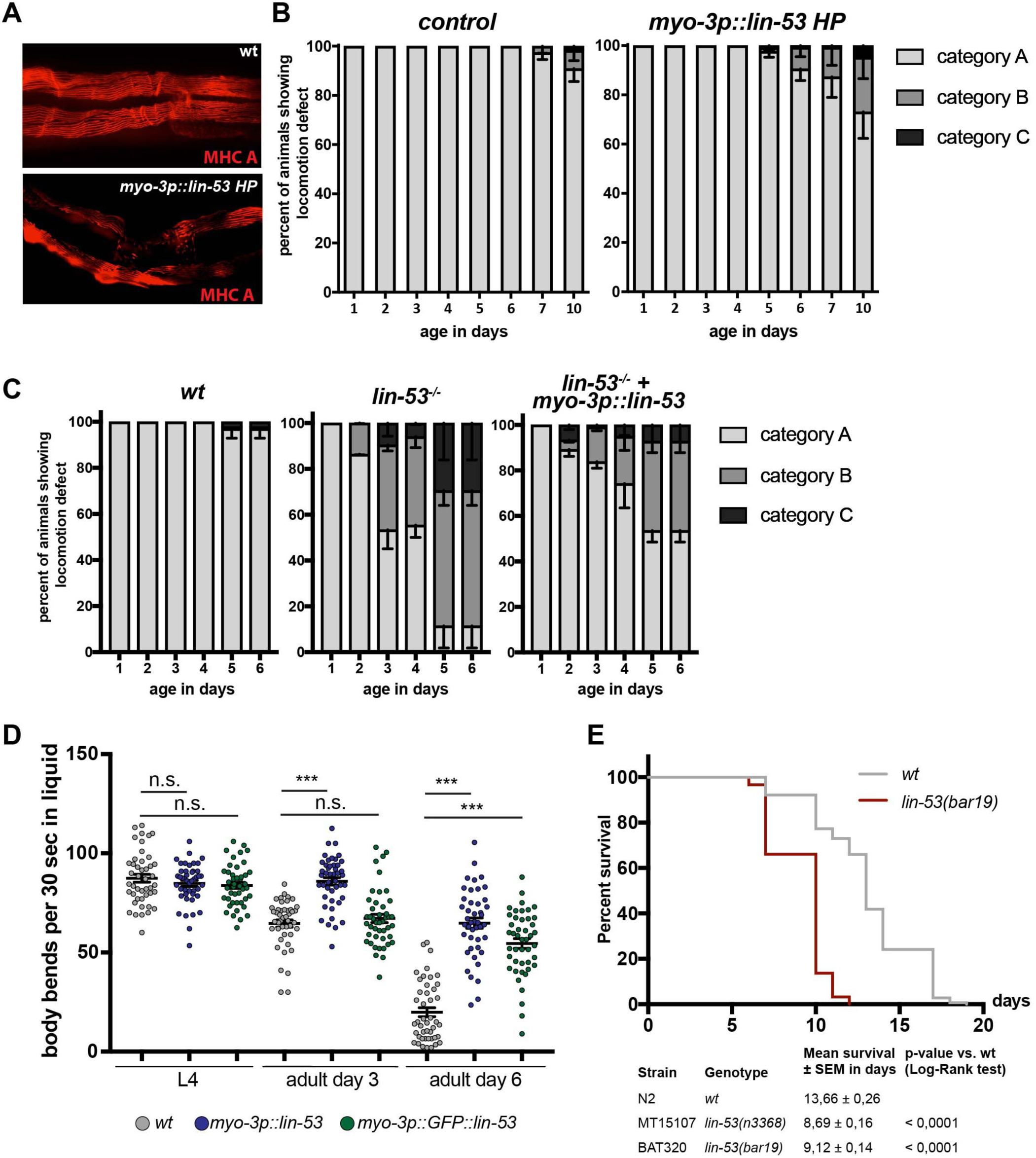
Loss of *lin-53* leads to motility defects due to disruption of muscles. Muscle-specific *lin-53* RNAi by expressing a *lin-53* hairpin construct only in the muscles analyzed using MHC immunostaining. The muscle structure is disrupted in *myo-3p::lin-53 HP* compared to control animals. (B) To analyze the motility of wt and *myo-3p::lin-53 HR* the assay was carried out on solid agar and motility was monitored throughout lifetime. Movement was classified in three different categories based on Herndon *et al.*, 2002 (Herndon et al., 2002): A = normal movement, B = impaired movement, C = no movement. Motility assays were carried out in triplicate using 50 animals per repeat. (C) Assay as in (B) with wt animals, *lin-53* mutants and *lin-53* mutants with muscle-specific *lin-53* overexpression (*myo-3p::lin-53*).. The impaired movement of *lin-53* mutants is rescued by *myo-3p::lin-53.* Motility assays were carried out using 50 animals per repeat. Scoring was started as animals reached day 1 of adulthood. (D) Overexpression of *lin-53* in muscles (*myo-3p::lin-53 and myo-3p::GFP::lin-53*) has beneficial effects on movement during aging. Body bends in liquid of L4 larvae, 3 days old adults, and 6 days old adults were measured. Upon overexpressing of *lin-53* in muscles at older stages of adulthood, worms move significantly better than control animals. Statistical analysis was carried out using unpaired one-way ANOVA; *** p < 0,0001, ns = not significant. (E) The CRISPR mutant *lin-53(bar19)* has a decreased lifespan by 4 days (p-value < 0,0001). Wild-type (wt) animals (grey line; mean lifespan 13,66 ± 0,26 days), *lin-53(bar19)* CRISPR mutants (red line; mean lifespan 9,12 ± 0,14 days), the experiment was carried out three times with at least 50 animals per repeat. Survival analysis was carried using Kaplan-Meier-estimator, p-value was calculated using Log-Rank Test.

**Fig. S2:**
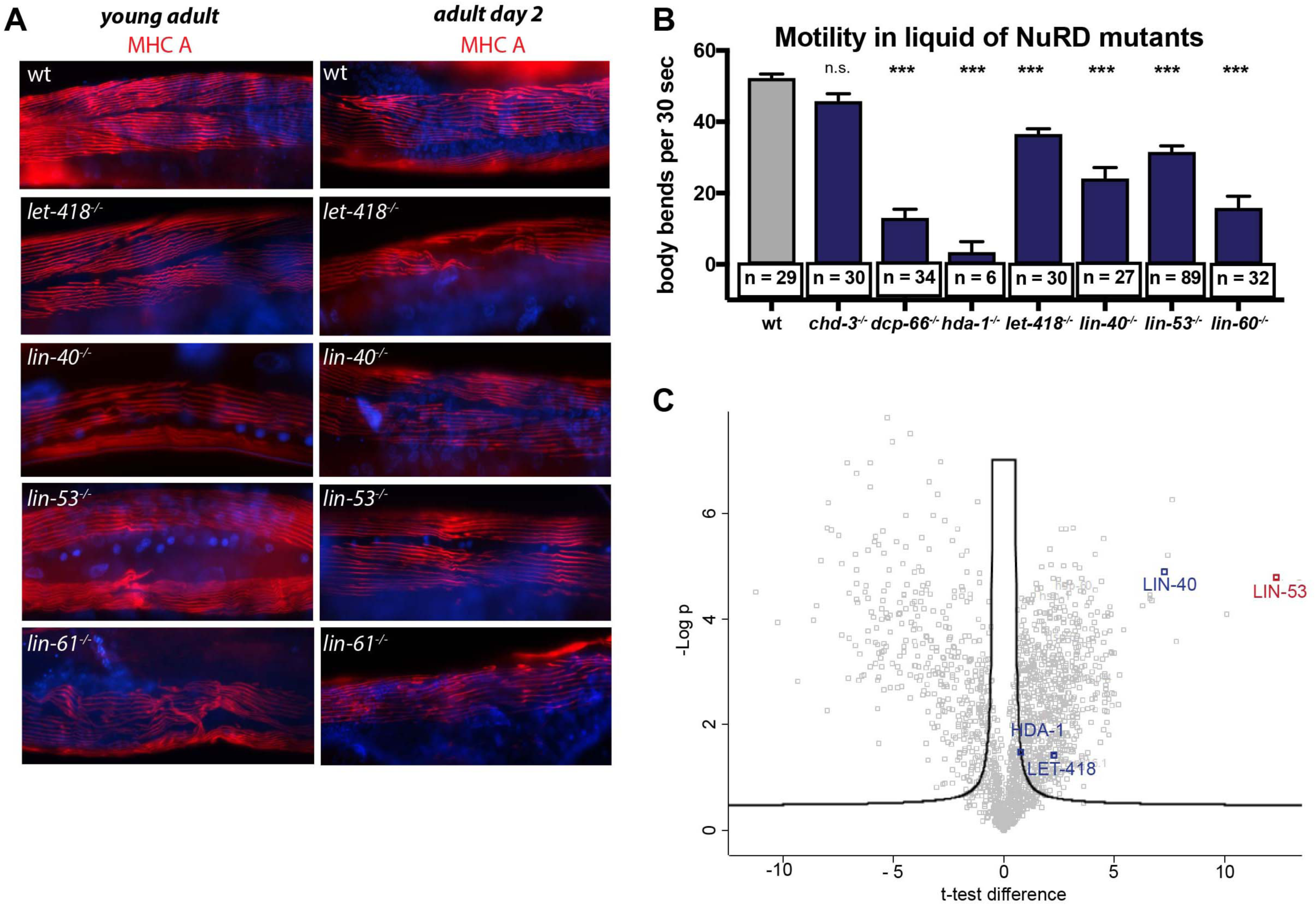
LIN-53 cooperates with NuRD to maintain muscle integrity. (A) Loss of the NuRD complex members disrupts muscle integrity as shown by immunostaining of myosin heavy chain (MHC) in *let-418, lin-40, lin-53* and *lin-61* mutants. Animals were at the young adult and 2 days old adult stage. (B) Motility of NuRD mutants in liquid. Thrashing assay was carried out as previously described. Statistics were done using One-way ANOVA (*** p < 0,0001). Number of animals counted is indicated below the columns. Animals were kept continuously at 25°C, before scoring at day 7 after hatching. (C) Volcano plot of IP-MS of muscle-specific LIN-53 (*myo-3p::lin-53::2xFLAG)*. Proteins enriched by LIN-53 pull-down were identified using permutation-based false discovery rate (FDR)-corrected two-sided t-test. The label-free quantification (LFQ) intensity of FLAG-pull down relative to control was calculated as difference and plotted on the x-axis against them −log_10_-transformed P-value of the t-test on the y-axis. The significance level (black line) was set to 0,05. Together with LIN-53 (red) the NuRD members LIN-40, LET-418 and HDA-1 (blue) were enriched suggesting that LIN-53 is interacting with the NuRD complex in muscles.

**Fig. S3:**
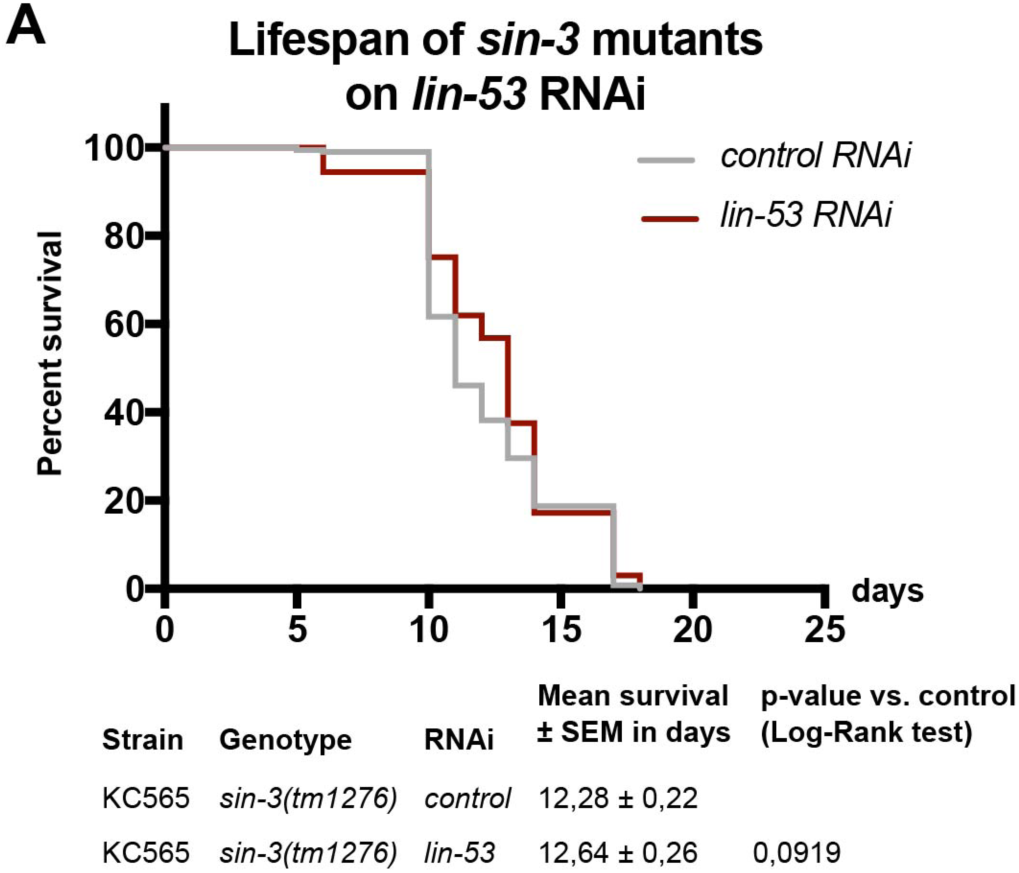
Depletion of *lin-53* in *sin-3* mutants does not further decrease lifespan. (A) Depletion of *lin-53* by RNAi in *sin-3* mutants does not change the lifespan compared to control RNAi (*lin-53* RNAi, red line; mean lifespan 12,64 ± 0,26 days; control RNAi grey line 12,28 ± 0,22 days; p-value < 0,0919). The experiment was carried out three times with 30 animals per repeat. Survival analysis was carried using Kaplan-Meier-estimator, p-value was calculated using Log-Rank Test.

**Fig. S4:**
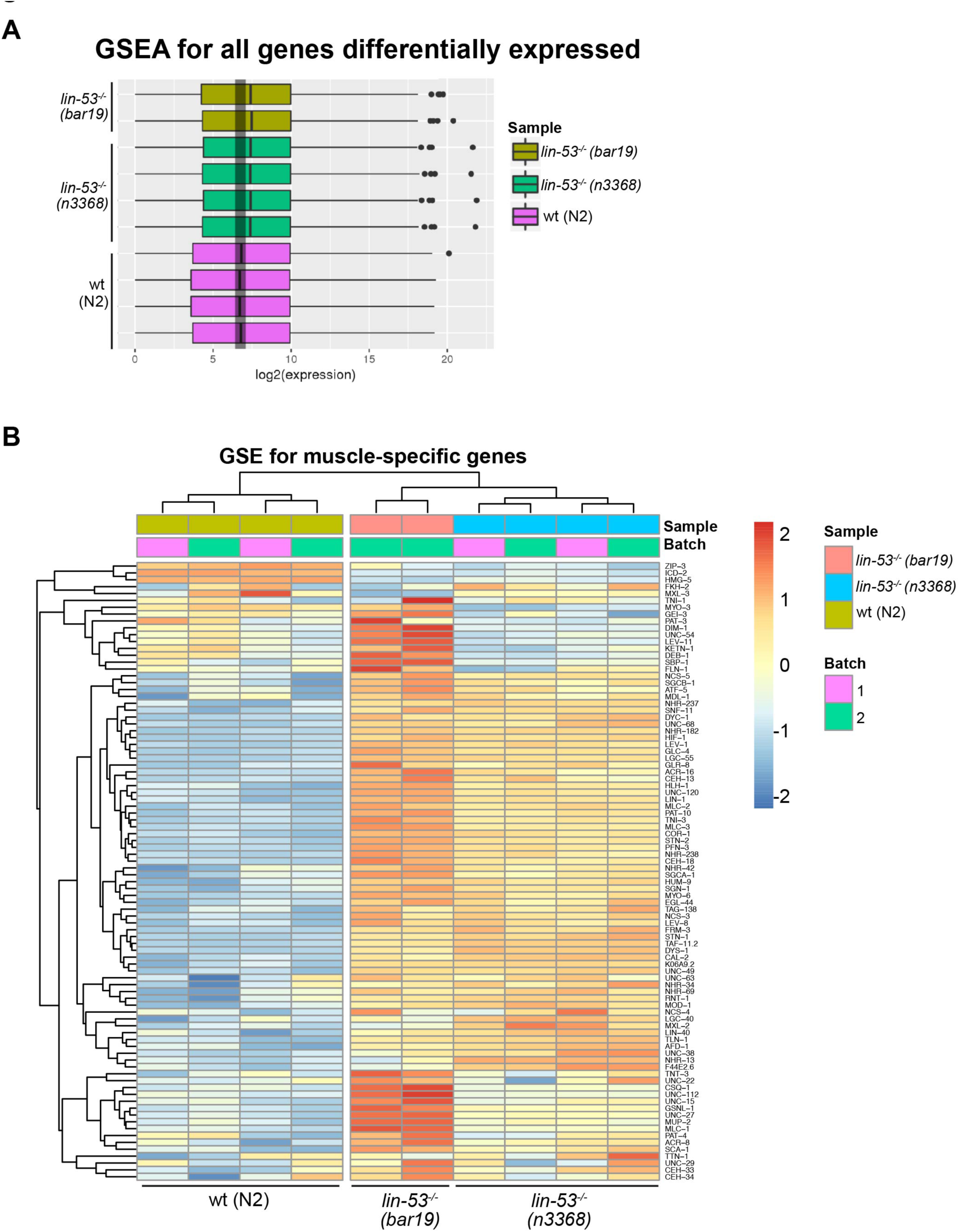
Transcriptome analysis of *lin-53* mutants. (A) Gene set enrichment analysis (GSEA) boxplots illustration for all gene families indicates that gene expression mostly increases upon loss of LIN-53. (B) Heat-map of the normalized expression values (VST) of muscle genes in *lin-53(n3368)* and *lin-53(bar19)* mutants compared to control animals.

**Fig. S5:**
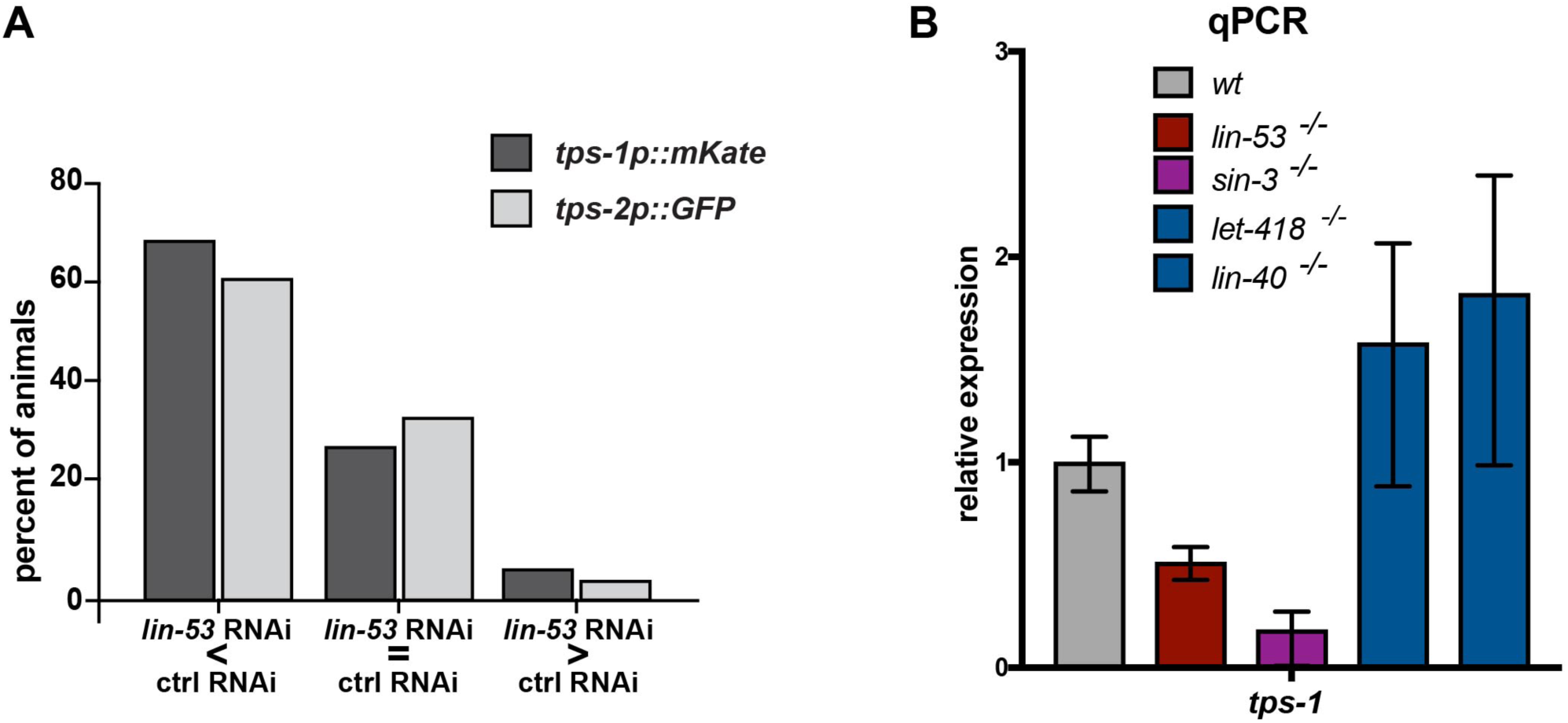
Loss of LIN-53 leads to decreased expression of Trehalose-phosphate synthase genes *tps-1* and *tps-2*. (A) Quantification of changed expression of *tps-1p::mKate* and *tps-2p::GFP* upon *lin-53* and control RNAi. Altogether 62 pairs of animals on *lin-53* and control animals were scored. More than 60 % of animals expressing the *tps-1p::mKate* or *tps-2p::GFP* reporter show decreased expression upon *lin-53* RNAi when compared to control animals. (B) qRT-PCR analysis of *tps-1* expression in *lin-53, sin-3*, *let-418*, and *lin-40* mutants. Expression decreases only in *lin-53* and *sin-3* mutants. Relative expression was calculated using the Livak method and statistics was done using unpaired t-test (*** p-value < 0,001, ** p-value < 0,05).

**Table S1:** Lifespan scoring data.

